# Iron deficiency affects early stages of embryonic hematopoiesis but not the endothelial to hematopoietic transition

**DOI:** 10.1101/462978

**Authors:** Maya Shvartsman, Saygin Bilican, Christophe Lancrin

## Abstract

Iron is an essential micronutrient for hematopoiesis and previous research suggested that iron deficiency in the pregnant female could cause anemia in the offspring. Since the development of all embryonic and adult blood cells begins in the embryo, we aimed to resolve the role of iron in embryonic hematopoiesis. For this purpose, we used an experimental system of mouse embryonic stem cells differentiation into embryonic hematopoietic progenitors. We modulated the iron status in cultures by adding either an iron chelator DFO for iron deficiency, or ferric ammonium citrate for iron excess, and followed the emergence of developing hematopoietic progenitors by flow cytometry. We found interestingly that iron deficiency by DFO did not block the endothelial to hematopoietic transition, the first step of hematopoiesis. However, it had a differential effect on the proliferation, survival and clonogenic capacity of hematopoietic progenitors. Surprisingly, iron deficiency affected erythro-myeloid Kit^pos^ CD41^+^ progenitors significantly more than the primitive erythroid Kit^neg^ CD41^+^. The Kit^pos^ progenitors paradoxically died more, proliferated less and had more reduction in colony formation than Kit^neg^ after 24 hours of DFO treatment. Kit^pos^ progenitors expressed less transferrin-receptor on the cell surface and had less labile iron compared to Kit^neg^, which could reduce their capacity to compete for scarce iron and survive iron deficiency. We suggest that iron deficiency could disturb hematopoiesis already at an early embryonic stage by compromising survival, proliferation and differentiation of definitive hematopoietic progenitors.

## Introduction

Embryonic hematopoiesis is an essential and complex process, which supplies blood to the developing embryo and the adult. It generates a wide range of hematopoietic progenitors and stem cells (HPSCs), from the primitive non-self-renewing erythroid progenitors of the yolk sac to the long-term self-renewing hematopoietic stem cells, which will reside in the adult bone marrow and generate all blood lineages when needed^1,2^. Despite its overall complexity, embryonic hematopoiesis can be simplified into a sequence of steps common for all HPSC types. It starts with the endothelial-to-hematopoietic transition (EHT), a step in which endothelial cells of a particular type (hemogenic endothelium) undergo through morphological and transcriptomic changes to become HPSCs^3-5^. Later on, these HPSCs proliferate, differentiate, and migrate to colonize their niches like the fetal liver, or the bone marrow^2, 6-8^. Recent work on human^9^ and mouse embryonic stem cells^10,11^ as well as on reprogrammed mouse embryonic fibroblasts^12^ contributed a lot to the knowledge of the transcription factors and the growth factors controlling embryonic hematopoiesis. Yet the role of iron in the steps of embryonic hematopoiesis is not completely understood.

Iron is an essential micronutrient required for catalysis, DNA synthesis, redox reactions and oxygen transport^13^. Iron deficiency through a knockout of iron import proteins like transferrin receptor (Tfrc) or Dmt1 (Slc11a2) causes anemia and embryonic or early postnatal lethality in mice^14,15^. Nutritional iron deficiency in the pregnant female increases the risk of iron deficiency and iron deficiency anemia in the offspring, according to animal models and human epidemiological studies ^16,17^. Hypotransferrinaemic hpx/hpx mice^18^ or mice chimeric for Tfrc knockout^19^ have a defect in T lymphoid differentiation, suggesting that the effects of iron deficiency might not be restricted only to the erythroid lineage. We therefore hypothesized that iron was important for an early step in embryonic hematopoiesis, which is common for all developing blood cells. Such a step could be either the EHT itself, or the steps right after it.

To dissect the step of embryonic hematopoiesis when iron is most required, we needed an experimental model where we could reproduce embryonic hematopoiesis, change the cellular iron status quickly and reversibly and test the effect of iron status on the steps of hematopoiesis in real time. To fulfill these requirements, we chose the experimental model of mouse embryonic stem cells progressively differentiating into blood through a hemangioblast stage^3^, similarly to what happens in yolk sac hematopoiesis^3^. The hemangioblast stage cultures start as Flk1^+^ mesoderm and differentiate with time into a mixed culture of endothelial, hematopoietic progenitor, and vascular smooth muscle cells^3^. In these cultures, we modified the cellular iron status by adding either an iron chelator (DFO) to cause iron deficiency or adding ferric ammonium citrate to cause iron excess^20^.

In this work, we demonstrated that iron deficiency by DFO did not block the EHT itself, but it differentially affected the proliferation, survival and differentiation of early hematopoietic progenitors. In contrast, iron excess had no adverse effects on hematopoietic progenitors. Thus, our findings offer broader understanding of how iron deficiency could affect embryonic hematopoiesis^16^.

## Results

### Iron deficiency by DFO does not inhibit EHT

We first tested whether iron deficiency would block the first step of embryonic hematopoiesis, which is the endothelial-to-hematopoietic transition (EHT)^3-5^. This hypothesis was based on previously published evidence that iron chelators were inhibiting the epithelial-to-mesenchymal transition (EMT), a mechanism similar to EHT^23^.

For this purpose, we differentiated mouse embryonic stem cells^21^ into hemangioblast cultures^3,10,11^ and followed the formation of hematopoietic progenitors from hemangioblast as a function of iron status and time. The time course of our experiments is schematically represented in Figure 1A. Our hemangioblast cultures were composed of four major cell types as demonstrated in Figure 1B: vascular smooth muscle cells (VSM), endothelial cells (EC), hematopoietic progenitor cells (HPCs) and cells in an intermediate EHT stage, referred to as Pre-HPCs. These four cell types were distinguished in flow cytometry by expression of an endothelial marker VE-Cadherin (VE-Cad) and an early hematopoietic marker CD41. Cells expressing neither VE-Cad nor CD41 were vascular smooth muscle cells; endothelial cells, VE-Cad^+^ CD41^-^; HPCs, VE-Cad^-^ CD41^+^ and Pre-HPCs, VE-Cad^+^ CD41^+^.

**Figure 1:**
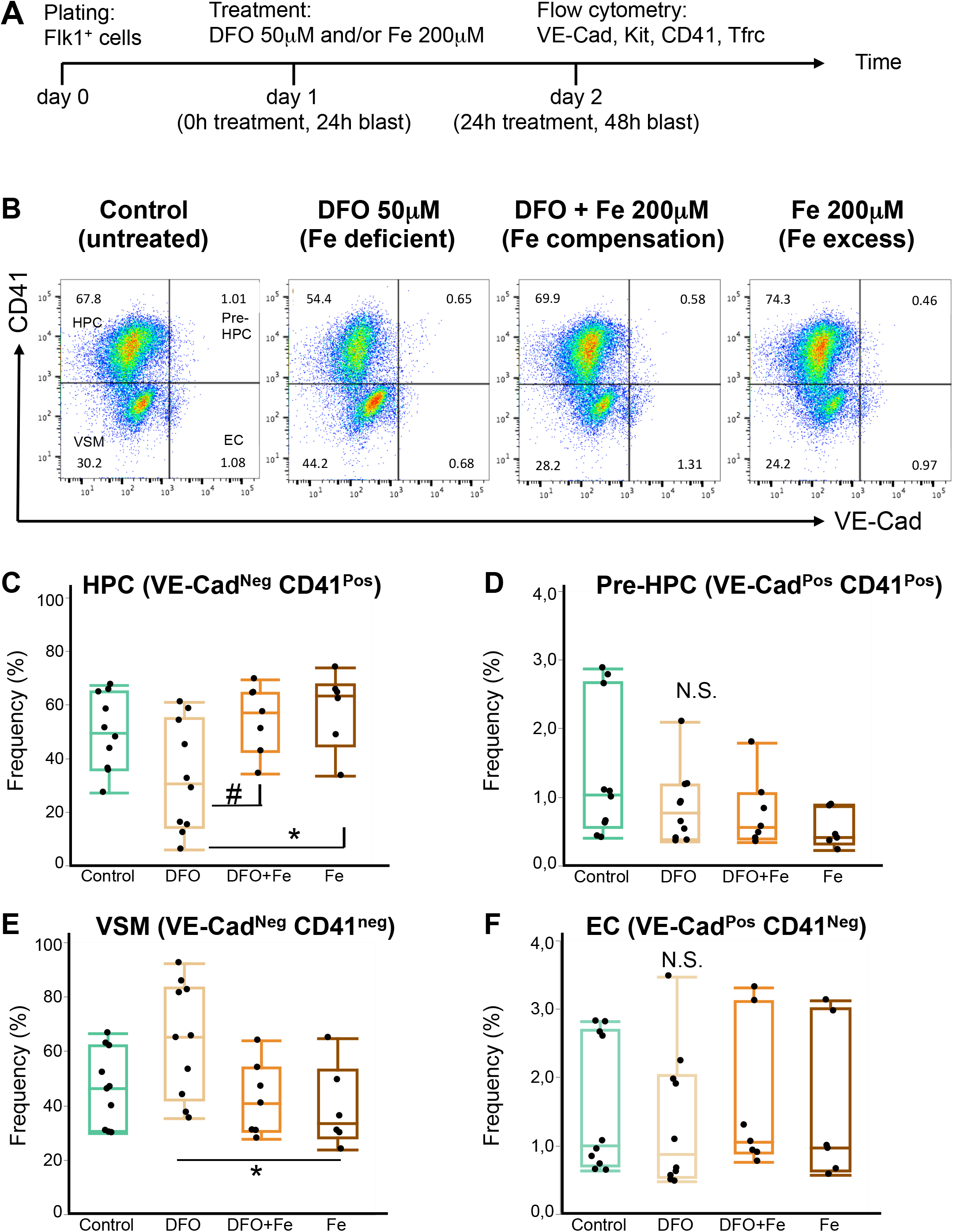
Iron deficiency does not block endothelial to hematopoietic transition. **(A)** A schematic timeline of an experiment in blast culture. Flk1^+^ cells were plated on day 0 in blast mix as previously described. 24 hours later the culture was treated with one of the following: nothing (control), DFO 50μM (iron-deficient conditions), DFO + ferric ammonium citrate 200μM (simultaneous neutralization of iron deficiency, or iron compensation), Fe 200μM (ferric ammonium citrate for iron excess conditions). After another 24 hours, all treatment groups were profiled with flow cytometry for expression of endothelial and hematopoietic cell surface markers. **(B)** A flow cytometry plot showing expression of CD41 against VE-Cad in day 2 blast (24 hours of treatment). According to the expression of these markers, each blast culture can be divided into 4 quadrants: VE-Cad^-^ CD41^-^ = vascular smooth muscle cells (VSM); VE-Cad^+^ CD41^-^ = endothelial cells (EC); VE-Cad^+^ CD41^+^ = pre-hematopoietic progenitor cells (Pre-HPC); VE-Cad^-^ CD41^+^ = hematopoietic progenitor cells (HPC). The experimental conditions are shown from left to right. **(C)** Tukey’s boxplots of HPC frequencies from flow cytometry experiments. Box whiskers show minimum and maximum, the line inside boxes shows median. For control and DFO groups n=10; for DFO+Fe and Fe groups n=6. All groups were analyzed by one-way ANOVA followed by Tukey’s test,. # p=0.049 and ^*^ p=0.027. **(D)** Tukey’s boxplots of Pre-HPC frequencies from flow cytometry experiments. Box whiskers show minimum and maximum, the line inside boxes shows median. For control and DFO groups n=10; for DFO+Fe and Fe groups n=6. All groups were analyzed by one-way ANOVA followed by Tukey’s test with p values shown on graphs. N.S. = not significant. **(E)** Tukey’s boxplots of VSM frequencies from flow cytometry experiments. Box whiskers show minimum and maximum, the line inside boxes shows median. For control and DFO groups n=10; for DFO+Fe and Fe groups n=6. All groups were analyzed by one-way ANOVA followed by Tukey’s test with p values shown on graphs. ^*^ Significant at p=0.032. **(F)** Tukey’s boxplots of EC frequencies from flow cytometry experiments. Box whiskers show minimum and maximum, the line inside boxes shows median. For control and DFO groups n=10; for DFO+Fe and Fe groups n=6. All groups were analyzed by one-way ANOVA followed by Tukey’s test with p values shown on graphs. N.S. = not significant.

In the case of an essential role of iron in EHT, we would expect that the addition of an iron chelator to our hemangioblast cultures would reduce HPCs frequency and cause accumulation of endothelial cells and/or Pre-HPCs^7^. DFO addition at 50 μM for 24 hours reduced the frequency and the absolute cell number of HPCs in culture (Figure 1B, 1C, Supplemental Figure 1). However, neither the frequency nor the cell number of endothelial and Pre-HPCs were increased by DFO (Figure 1B, 1D, 1F, Supplemental Figure 1), suggesting no accumulation of these cells took place. We further observed an increase in the frequency of vascular smooth muscle cells after DFO treatment (Figure 1E), but their absolute cell number was not increased (Supplemental Figure 1).

Excess iron added as 200 μM ferric ammonium citrate together with DFO abrogated all effects of DFO in culture, demonstrating that the observed effects were truly due to iron deficiency. Iron addition together with DFO or alone did not reduce the frequencies and absolute cell numbers of any cell type compared to untreated control (Figure 1, Supplemental Figure 1), suggesting that in our conditions iron excess was at least not toxic to cells.

### Iron deficiency differentially affects hematopoietic progenitors

All hematopoietic progenitors in our cultures can be divided into two major subtypes: definitive Kit^pos^ HPCs (Kit^+^ CD41^+^) and primitive Kit^neg^ HPCs (Kit^-^ CD41^+^). The Kit^pos^ HPCs mostly yield multilineage hematopoietic colonies when plated on methylcellulose with appropriate growth factors and are capable to reconstitute irradiated mice albeit transiently, while the Kit^neg^ HPCs yield mostly primitive erythroid colonies on methylcellulose and have no reconstitution capacity^3,24^. When we examined the effect of DFO on both kinds of HPCs, we saw that the frequency of Kit^pos^ HPCs was significantly (p<0.0001) reduced compared to untreated control (Figure 2A,B) and the frequency of Kit^neg^ HPCs was not significantly changed. The cell numbers of both progenitor types were significantly reduced, but the decrease in Kit^pos^ HPCs was stronger (Supplemental Figure 2). This observation was surprising, since we did not expect to find a differential effect of iron deficiency on hematopoietic progenitors at such an early stage. Endothelial and vascular smooth muscle cells, viewed in these experiments as Kit^+^ CD41^-^ and Kit^-^ CD41^-^ respectively, behaved similarly to the experiments described in Figure 1 (Figure 2 and Supplemental Figure 2).

**Figure 2:**
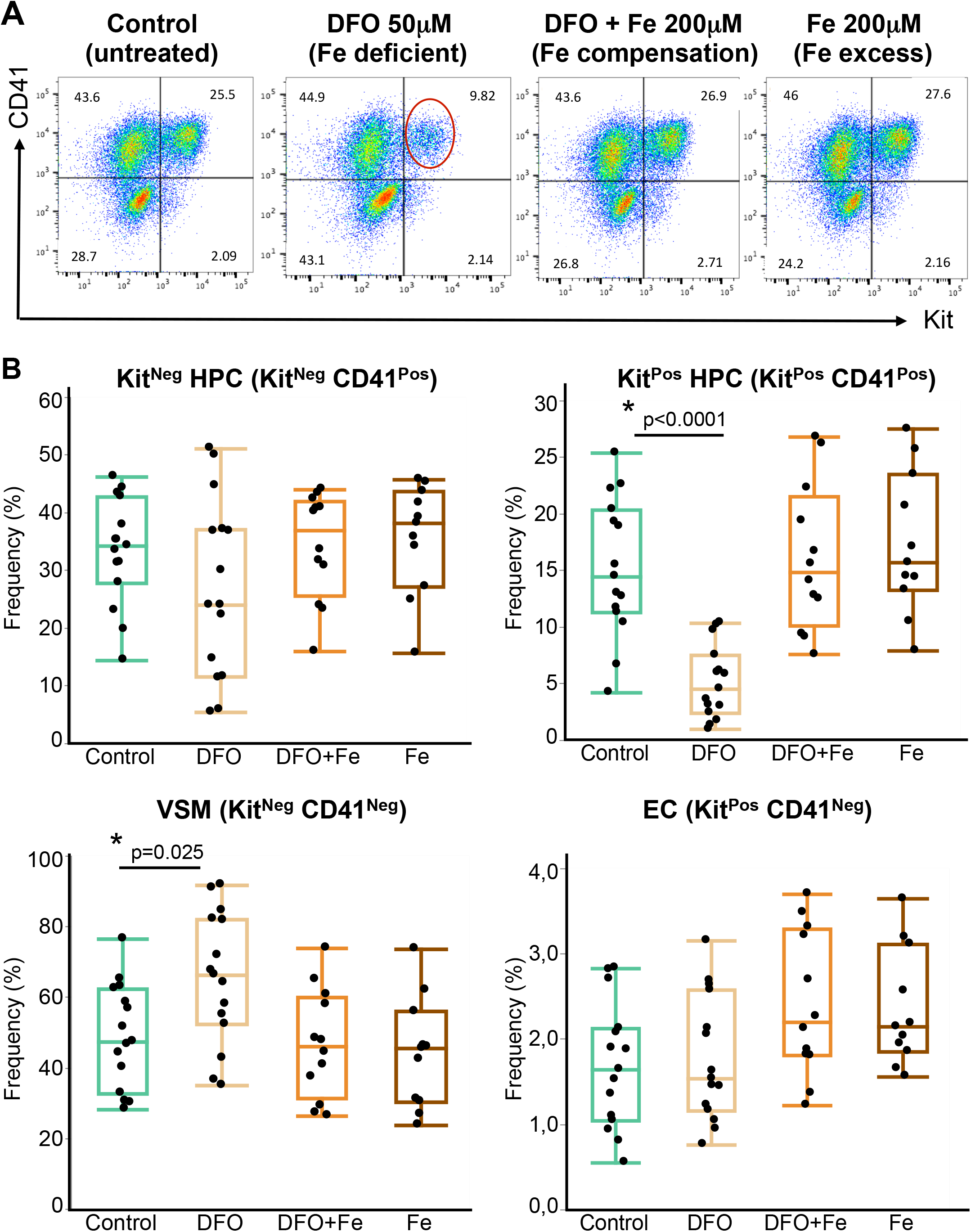
Iron deficiency reduces the frequency of Kit^Positive^ Hematopoietic Progenitor Cells. **(A)** A flow cytometry plot showing expression of CD41 against Kit in day 2 blast (24 hours of treatment). According to the expression of these markers, each blast culture can be divided into 4 quadrants: Kit^-^ CD41^-^ = VSM; Kit^+^ CD41^-^ = EC; Kit^+^ CD41^+^ = Kit positive HPC (Kit^Pos^ HPC); Kit^-^ CD41^+^ = cKit negative HPC (Kit^Neg^ HPC). The experimental conditions are shown from left to right. The red circle shows the Kit^Pos^ HPC population whose frequency decreased following the DFO treatment. **(B)** Tukey’s boxplots of cell type frequencies from flow cytometry experiments. Box whiskers show minimum and maximum, the line inside boxes shows median. For control and DFO groups n=15; for DFO+Fe group n=13 and Fe group n=11. All groups were analyzed by one-way ANOVA followed by Tukey’s test with p values shown on graphs.

### Iron deficiency does not affect hematopoietic progenitor identity

To find an explanation for the observed differential effect of iron deficiency on HPCs, we profiled gene expression in both kinds of progenitors by single-cell quantitative RT-PCR (sc-q-RT-PCR). We isolated single cells from Kit^Neg^ HPCs, Kit^Pos^ HPCs, VSM and ECs and tested a panel of 96 genes, among them endothelial (*Cdh5*, *Ramp2*, *Sox17* and *Erg*), hematopoietic (erythroid and myeloid), smooth muscle (*Acta2*, *Col3a1*, *Pdgfrb*, *Meis2* and *Snai1*) and iron metabolism (*Fth1*, *Tfrc* and *Slc11a2*) genes (Supplemental Figure 3). Interestingly, the four populations defined by Kit and CD41 cell surface markers segregated into four distinct cell clusters following hierarchical clustering and Principal Component Analysis (PCA) analysis (Fig. 3 and Supplemental Figure 3). The Kit^Neg^ HPCs were clearly erythroid, expressing genes of embryonic hemoglobins (*Hbb-y*, *Hbb-bh1* and *Hba-x*), *Epor*, *Gata1* and *Aqp8*. In contrast, the Kit^Pos^ HPCs were composed of cells expressing erythroid genes (albeit at a lower level than Kit^Neg^ HPCs) and others expressing white blood specific genes (*Spi1*, *Alox5ap*, *Coro1a* and *Mpo*). All HPCs and non-hematopoietic cells highly expressed the *Fth1* gene, which encodes one of the ferritin protein subunits.

**Figure 3:**
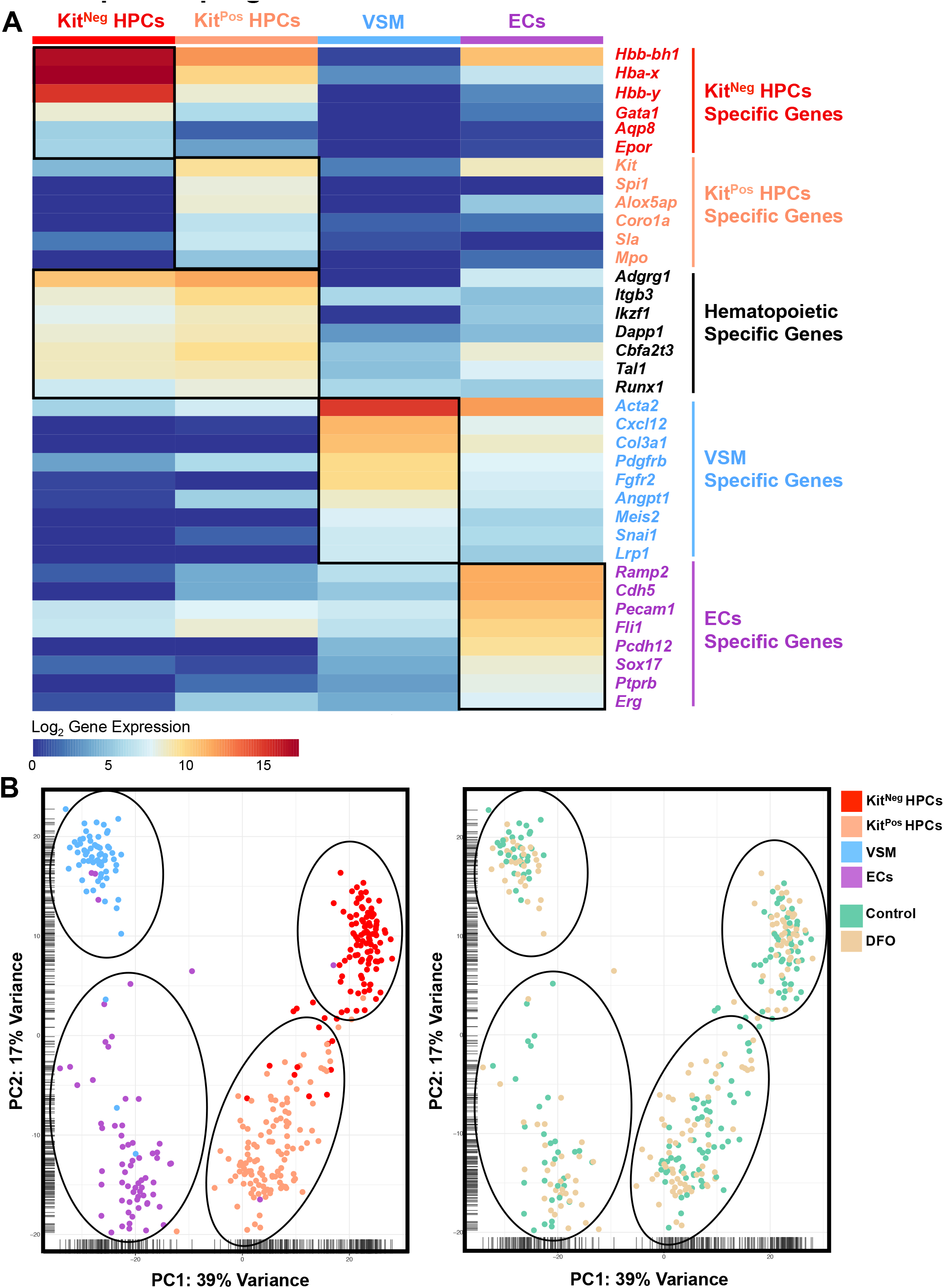
Iron deficiency does not change the identity of hematopoietic progenitors. **(A)** Heatmap showing the expression of key genes for the indicated populations found in control condition. The four groups were based on the phenotype of sorted single cells. **(B)** PCA plots showing the 366 cells tested by sc-q-RT-PCR (Control and DFO). The PCA plot on the left shows the four major cell clusters and the PCA on the right shows the distribution of cells from Control and DFO experimental conditions.

Remarkably, DFO-treated cells of any cell type clustered together with its respective control cells, suggesting that iron deficiency had no large-scale effect on the expression profile of the selected gene panel as shown by hierarchical clustering, PCA and Anova pairwise analysis (Figure 3, Supplemental Figures 3 and 4).

### Iron deficiency differentially affects proliferation and survival of hematopoietic progenitors

Since iron deficiency by DFO selectively reduced the frequency of Kit^Pos^ HPCs, we investigated the causes of this reduction. Since the EHT is not inhibited by the DFO treatment (Figure 1), we hypothesized that the Kit^Pos^ HPCs frequency could be reduced because of a decrease in proliferation or an increase in cell death. We measured cell proliferation in control, iron deficient and iron-excess conditions with the aid of a ClickIt-EdU kit, which only labels cells in S phase through incorporation of EdU^11^. Overall, hematopoietic progenitors were the most proliferating cells in culture having between 54-64% of S-phase cells, while endothelial and vascular smooth muscle cells proliferated less, with 23-30% (Figure 4 and Table 2). In control conditions, both Kit^Pos^ and Kit^Neg^ HPCs had similar proliferation rate, but the decrease in proliferation after DFO was significantly stronger in Kit^Pos^ HPCs. Adding excess iron together with DFO kept cell proliferation to control levels.

**Figure 4:**
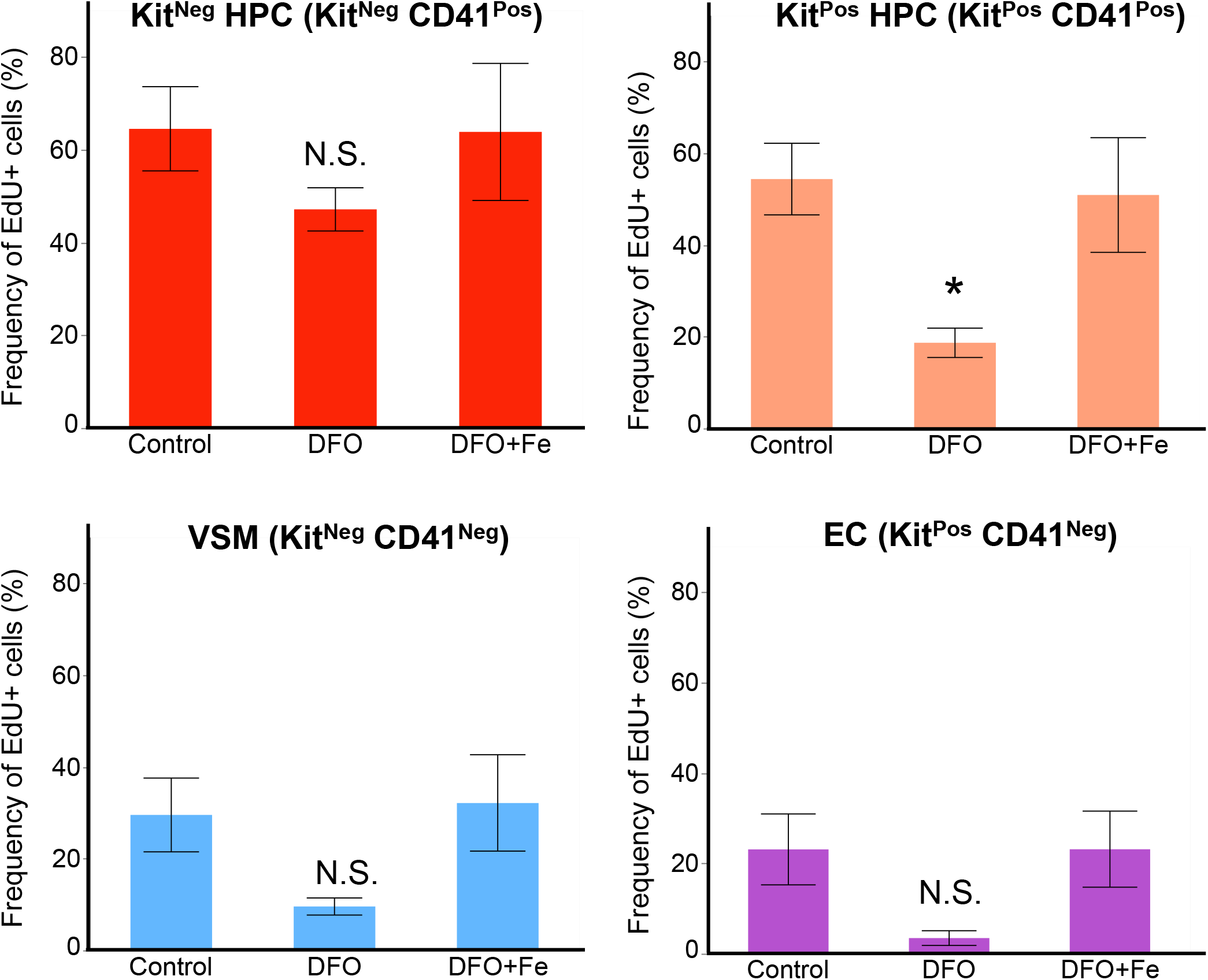
Iron deficiency reduces the proliferation of Kit^Pos^ HPCs. The frequency of EdU^+^ cells (i.e. in S-phase) is shown as a function of treatment for each of the cell types in the blast culture. Data are shown as mean ± SE, n=4. ^*^ p<0.05, one-way ANOVA and Tukey’s multiple comparisons test.

Our apoptosis measurements using AnnexinV and 7AAD^22^ demonstrated that in control conditions the death rate of both progenitor types was not significantly different (Figure 5A,B and Table 1). DFO treatment increased the frequency of late apoptotic cells in both HPC types compared to control or iron-treated groups (Figure 5A). This increase in AnnexinV^+^ 7AAD^+^ cells was seen at 24 and 48 hours of DFO 50 μM treatment. Excess iron added together with DFO displayed apoptotic cell frequencies close to control levels. Since there was some extent of cell death in control conditions, we calculated the net cell death as delta apoptosis by subtracting the frequency of apoptotic cells in control conditions from the frequency of apoptotic cells with DFO (Δ _DFO – control_). The net cell death was significantly higher in Kit^pos^ HPCs than in Kit^neg^ HPCs in all time-points of DFO treatment (Figure 5B). Together, our results suggest that DFO reduces Kit^Pos^ HPCs frequency both by reducing proliferation and by increasing cell death.

**Figure 5:**
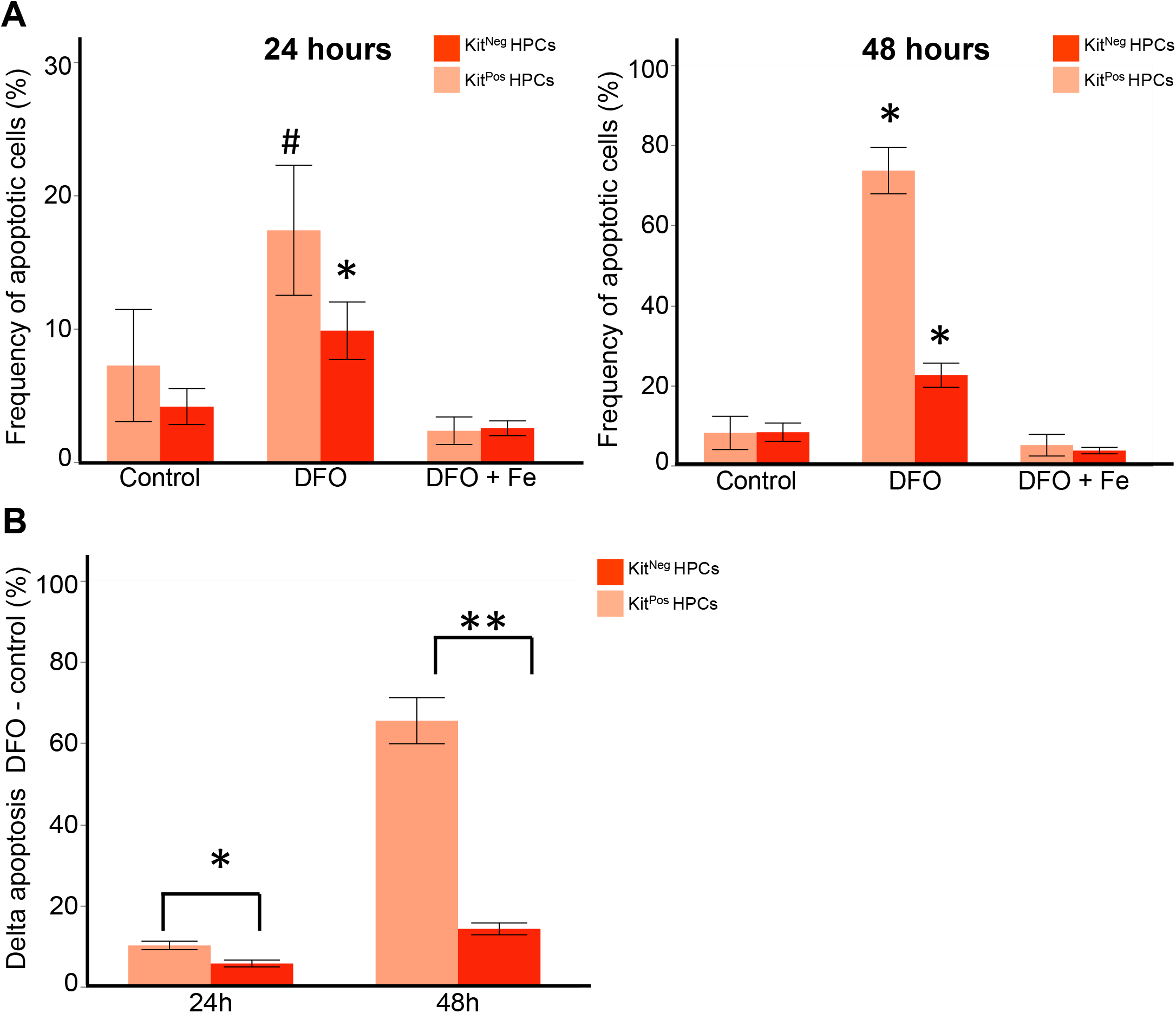
Iron deficiency leads to a higher apoptosis rate of KitPos HPCs compared to KitNeg ones. **(A)** The frequency (%) of apoptotic (AnnexinV^+^ 7AAD^+^) cells is presented as a function of treatment in both Kit^Pos^ and Kit^Neg^ HPCs after 24 hours (left panel) and 48 hours (right panel) treatment. *p<0.05 for control versus DFO or # for DFO versus Fe groups by one-way ANOVA and Tukey multiple comparisons test. **(B)** The delta apoptosis (% DFO - % control) was compared between Kit^+^ CD41^+^ and Kit^-^ CD41^+^ HPs. *p<0.05 and **p<0.01 for Kit^+^ CD41^+^ versus Kit^-^ CD41^+^ by paired two-tailed t-test.

**Table 1:**
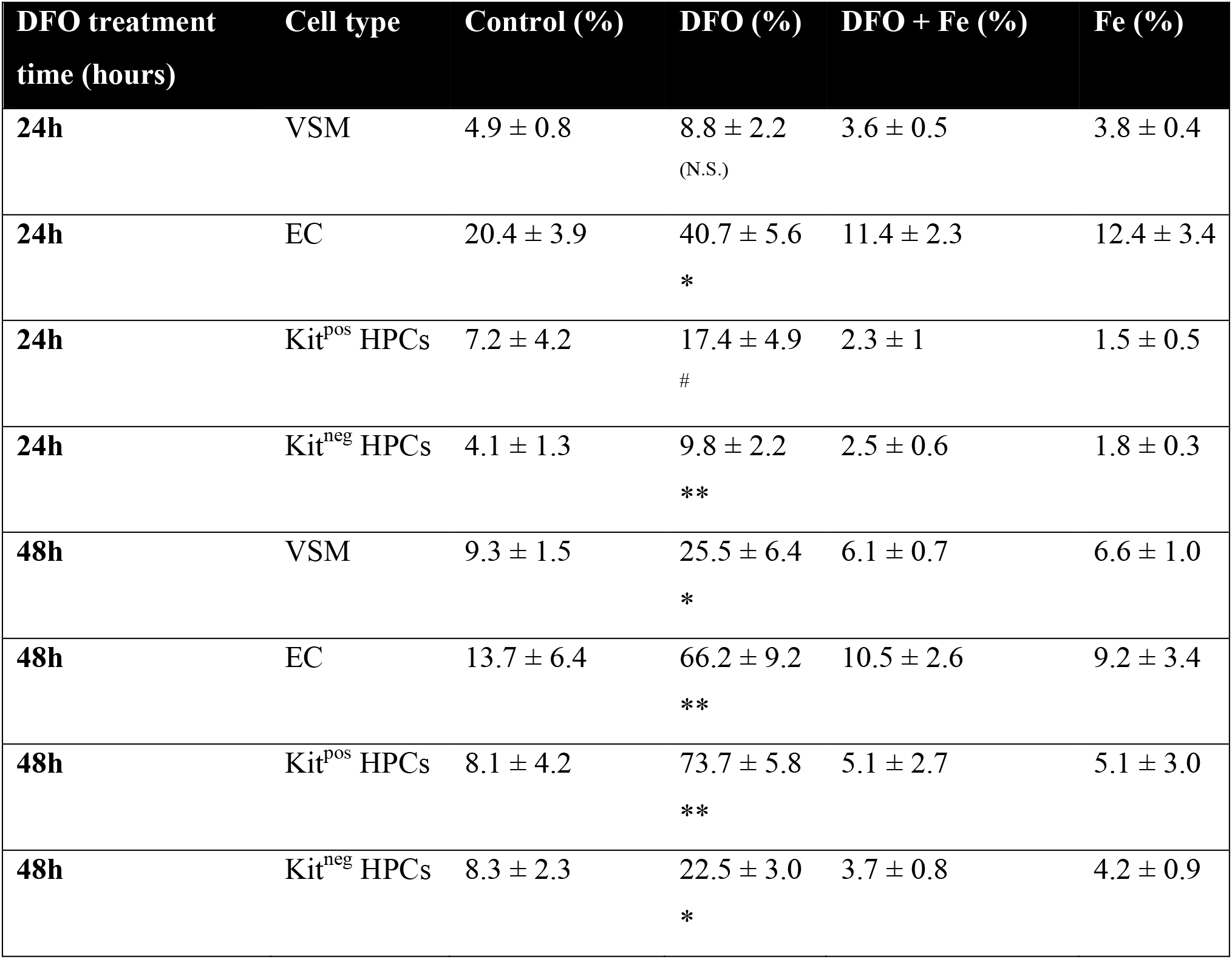
Iron deficiency differentially increases cell death. Cell death was measured by flow cytometry as a frequency (%) of AnnexinV+ 7AAD+ cells from each cell type. Data are presented as mean ± SE, N=4. N.S. – not significant DFO vs control ^*^ Significant DFO vs control at p<0.05, one-way ANOVA + Tukey ^**^ Significant DFO vs control at p<0.01, one-way ANOVA + Tukey ^#^ Significant DFO vs Fe treated groups, one-way ANOVA + Tukey

**Table 2.**
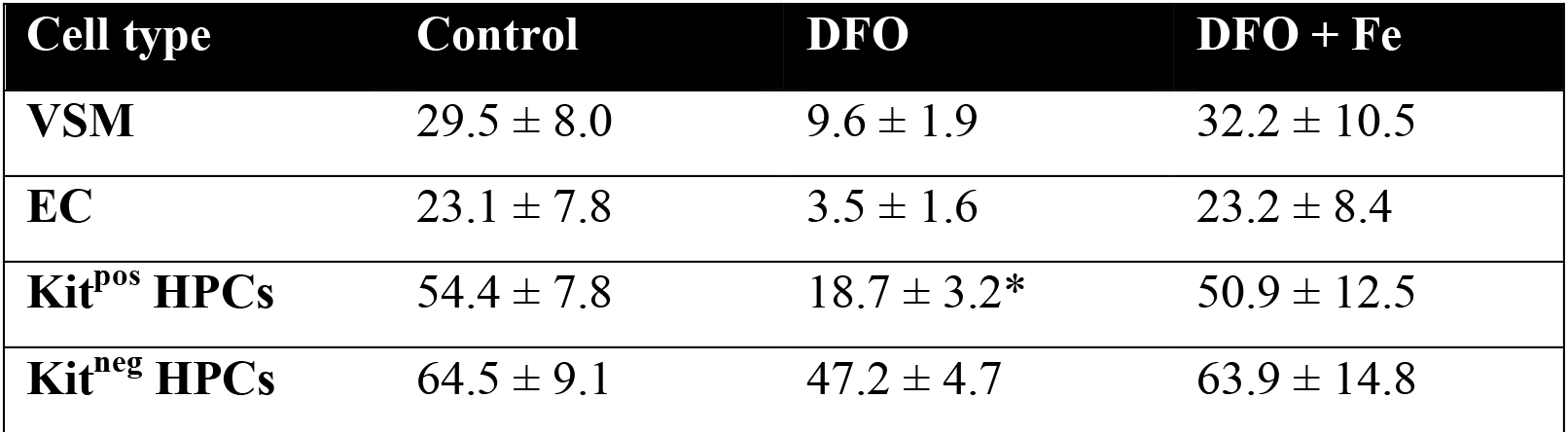
Iron deficiency reduces proliferation of Kit^pos^ HPCs. Cell proliferation was measured as a frequency (%) of S-phase EdU-positive cells. Data are presented as mean ± SE, N=4 ^*^ Significant DFO vs control at p<0.05, one-way ANOVA + Tukey

### Iron deficiency reduces colony-forming capacity of both Kit^-^ and Kit^+^ hematopoietic progenitors

We aimed to resolve whether iron deficiency would affect the differentiation of early hematopoietic progenitors into mature blood lineages. For this purpose, we sorted both kinds of HPs from control and DFO-treated conditions and put them on DFO-free methylcellulose plates for a week. As published previously^24^ and in correspondence with our above-mentioned sc-q-RT-PCR results, Kit^neg^ HPCs gave mostly primitive erythroid colonies after one week on methylcellulose; the Kit^pos^ HPCs gave rise to erythroid, erythromyeloid and macrophage colonies. The clonogenic capacity of Kit^pos^ HPCs was higher than of Kit^neg^, in agreement with previously published work^3,24^. DFO treatment significantly reduced the total amount of colonies formed from all HPCs together (Figure 6) and the amount of erythroid and macrophage colonies from Kit^pos^ HPCs. We did not observe a statistically significant effect of DFO on colony output from Kit^neg^ HPCs. Overall, DFO treatment of early HPCs for 24 hours was sufficient to compromise HP clonogenic capacity, disturbing differentiation in the longer term.

**Figure 6:**
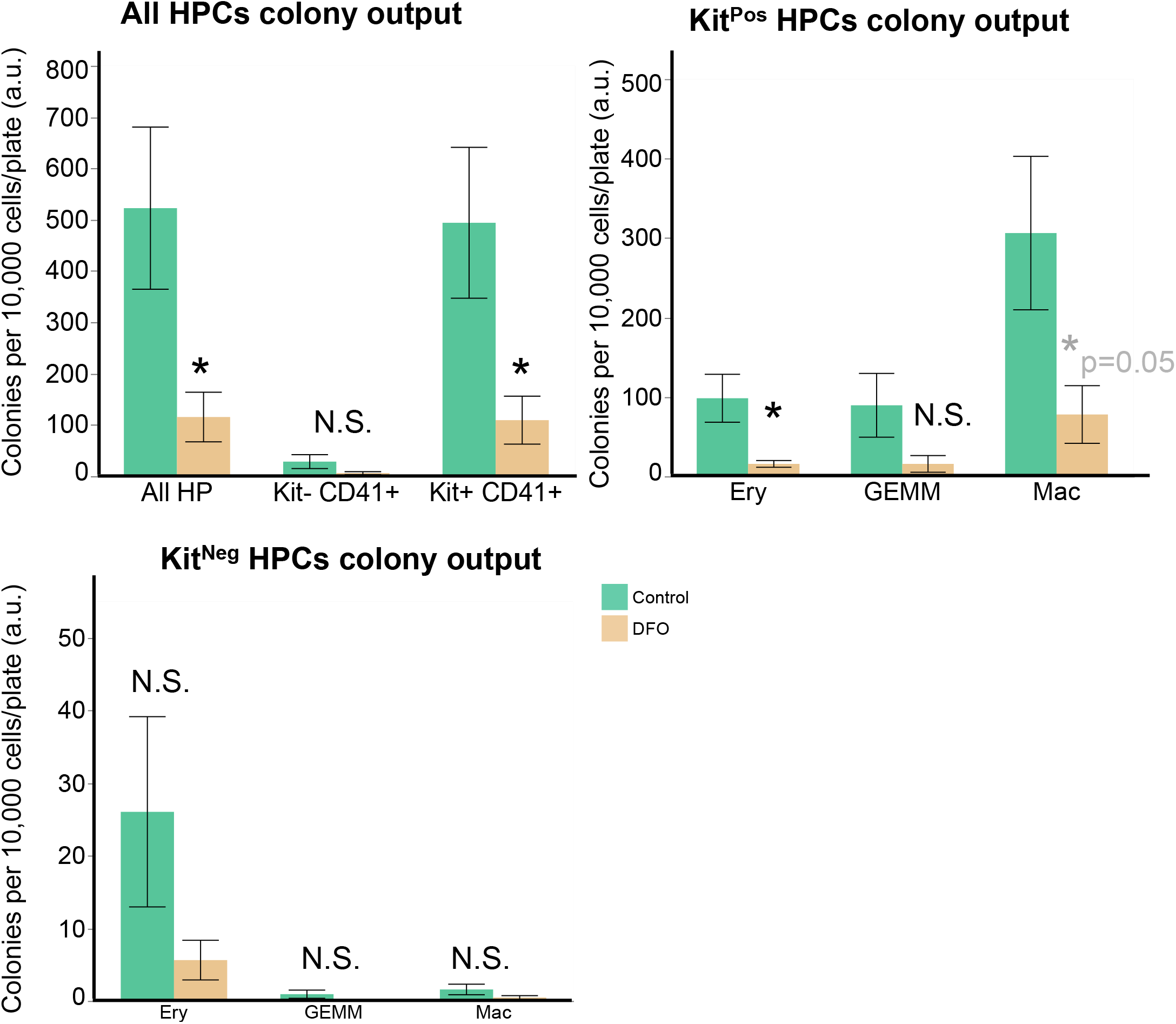
Iron deficiency reduces hematopoietic colonies’ formation. Colony formation in HPs isolated from control or DFO 50μM-treated day 2 blast cultures. Data are presented as mean ± SE, n=5. Colonies were calculated per 10,000 cells/plate for all HPCs; Kit^Pos^ HPCs; Kit^Neg^ HPCs. *p<0.05, paired two-tailed t-test; N.S. = non-significant control versus DFO.

### Hematopoietic progenitors differ in their endogenous transferrin receptor protein expression and labile iron levels

We hypothesized that the hematopoietic progenitors the most sensitive to iron deficiency should have higher metabolic demand for iron, reflected by higher expression of transferrin receptor (Tfrc) protein on the cell surface. However, our flow cytometry studies paradoxically showed the opposite. The more sensitive Kit^Pos^ HPCs had significantly lower Tfrc expression than Kit^neg^ HPCs, as seen by both the frequency (%) of Tfrc^+^ cells and the mean Tfrc fluorescence (Figure 7 and Table 3). Overall in culture, hematopoietic progenitors had higher Tfrc expression than non-hematopoietic cells, and most of the Tfrc fluorescence came from Kit^Neg^ HPCs. Iron deficiency or iron excess gave a trend of increase or decrease of Tfrc expression respectively, but the Tfrc level was always higher in Kit^Neg^ HPCs compared to other cell types (Table 3). The Kit^Neg^ HPCs also had the highest labile iron levels in culture as detected by cytosolic calcein (Table 3). Together, these observations could explain our paradox of HPCs sensitivity to iron deficiency: the more sensitive Kit^Pos^ HPCs express less Tfrc and have less labile iron inside, therefore are less able to compete for scarce iron and survive through iron deficiency.

**Table 3:** Kit^neg^ HPCs have highest Tfrc and labile iron compared to other cell types. Transferrin receptor levels on the cell surface were measured by flow cytometry as described in Methods. Data are presented as mean ± SE, N=4. Cytosolic Labile iron levels in control untreated cells were measured with calcein by flow cytometry as described^20^. Data are presented as mean± SE, N=3. ^*^ Significant in Kit^Neg^ HPCs vs other cell types.

| Cell type | Mean Tfrc fluorescence (a.u.) |  |  | Basal labile iron (r.u.) |
| --- | --- | --- | --- | --- |
|  | Control | DFO | DFO + Fe |  |
| VSM | 672 ± 152 | 786 ± 183 | 485 ± 80 | 0.09 ± 0.02 |
| EC | 1026 ± 377 | 880 ± 130 | 429 ± 113 | 0.22 ± 0.01 |
| Kit<sup>pos</sup> HPCs | 2253 ± 699 | 2529 ± 423 | 634 ± 124 | 0.20 ± 0.02 |
| Kit<sup>neg</sup> HPCs | 11301 ± 2841* | 12889 ± 2301* | 2325 ± 438* | 0.45 ± 0.1 * |

## Discussion

In the present work, we aimed to resolve the role of iron in early hematopoiesis. While previous studies^14^,^15^ demonstrated that iron deficiency causes anemia, some other works in Tfrc^-/-^ chimeric mice or the atransferrinemic mice showed that iron deficiency could also affect non-erythroid lineages^18-19^. Even in the total Tfrc knockout mouse, which had anemia and embryonic lethality by E12.5 at the latest^14^, there still was some yolk-sac erythropoiesis, suggesting that some hematopoietic progenitors and stem cells could be more sensitive to iron deficiency than others. We therefore hypothesized that iron deficiency could affect not only erythroid development, but rather affect a hematopoietic step common for all blood lineages.

We used an *in vitro* model system of mouse embryonic hematopoiesis, which allowed us to visualize the early steps of the process in real-time^3,10,11^. It has the advantages of accessibly and reversibly modifying the cellular iron by chemical/pharmacological means, and lacking complications like embryonic lethality.

The addition of DFO to our hemangioblast cultures did not cause a massive accumulation of endothelial or Pre-HPCs characteristic of TGFβ treatment or overexpression of key transcription factors^10,11^. Neither the frequency nor the cell number of endothelial cells were increased (Figures 1,2 and Supplemental Figures 1, 2). Therefore, we conclude that DFO does not block the EHT per se.

In our cultures, we observed two kinds of hematopoietic progenitors, which differ by their Kit expression Kit^Pos^ and Kit^Neg^ HPCs. DFO had an unexpected differential effect on HPCs, reducing the frequency and cell number of Kit^Pos^ more than of Kit^Neg^ (Figure 2). These early progenitors differed in their endogenous transcriptomic profile (Figure 3) already at this early stage. The Kit^Pos^ HPCs were erythro-myeloid, while the Kit^Neg^ were primitive erythroid by their gene expression profile. That difference in gene expression was also consistent with the HPC differentiation profile: Kit^pos^ HPCs gave mixed lineage colonies after a week of CFU assay (Figure 6), and Kit^neg^ HPCs gave mostly primitive erythroid colonies. Our observations were consistent with previously published works^3,10,11,24^. DFO treatment did not change the overall transcriptomic profile of any HPCs (Figure 3) and it did not change the type of colonies generated on methylcellulose (Figure 6), suggesting that iron deficiency does not affect cell identity or differentiation direction. The overall HPC differentiation capacity, reflected by colony number, was reduced by DFO (Figure 6) presumably due to inhibition of proliferation and increased cell death (Figures 4,5 and Tables 1,2). While the effects on proliferation and cell death were apparent right after completion of 24 or 48-hours treatment, the consequences on differentiation were long lasting. We could therefore suggest that iron deficiency could have both short and long-term effects on hematopoiesis.

We were interested in the reason for the differential sensitivity of our HPCs to DFO. In general, the sensitivity of a cell to iron deficiency could be determined by a combination of factors: the iron-import capacity of a cell; the iron content inside the cell; and the metabolic requirement for iron. We initially assumed that the more sensitive progenitors would be more erythroid, requiring more iron for hemoglobin synthesis; and would express more transferrin receptor, but we actually observed the opposite. We unexpectedly found that the less erythroid Kit^Pos^ HPCs were the more sensitive ones. When we measured the Tfrc and iron levels in both HPC types, we found that more sensitive Kit^Pos^ HPCs had endogenously lower Tfrc expression and less intracellular iron than the Kit^Neg^ HPCs (Figure 7 and Table 3). Having less Tfrc on the cell surface, the Kit^Pos^ HPCs could be less capable to compete for scarce iron with Kit^Neg^. Having less iron inside, the Kit^Pos^ HPCs could be less capable to survive iron deficiency.

**Figure 7:**
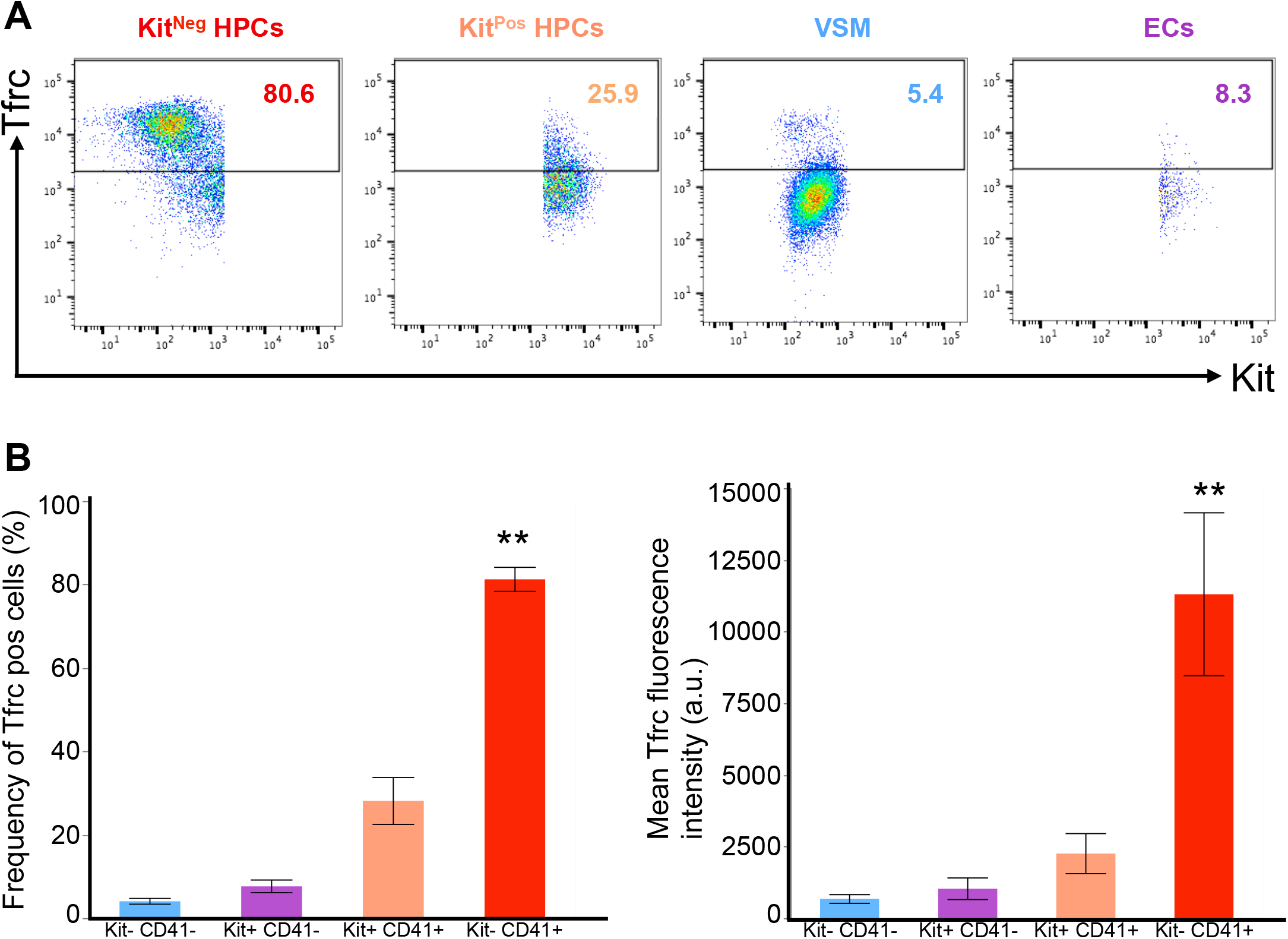
Kit^Neg^ HPCs express the highest level of Transferrin receptor. **(A)** Representative flow cytometry plots showing Tfrc (CD71) versus Kit expression in different cell types. **(B)** Frequency (%) of cells being positive for Tfrc in different cell types. Data are shown as mean ± SE. ** Significant in Kit^Neg^ HPCs versus Kit^Pos^ HPCs at p<0.01, ANOVA and Tukey-Kramer. **(C)** Mean Tfrc fluorescence in arbitrary units in different cell types. ** Significant in Kit^Neg^ HPCs versus Kit^Pos^ HPCs at p<0.01, ANOVA and Tukey-Kramer.

Overall, we demonstrate for the first time that iron deficiency could affect hematopoiesis at an unexpectedly early embryonic stage. It is known from previous works that iron deficiency in pregnant rats reduced the total iron content in whole embryos^16^ and fetal liver^25^, as well as hematocrit, hemoglobin and RBC counts in the offspring^16^. Our study suggests that iron deficiency could have consequences beyond erythropoiesis, also affecting developing myeloid cells in the embryo. Our work indicates that other aspects of hematopoiesis besides erythropoiesis need also to be assessed in cases of iron deficiency.

## Materials and Methods

### Maintenance and differentiaton of mouse embryonic stem cells (mESCs)

We used the A2lox Cre mESC cell line, which was a kind gift from Michael Kyba^21^. The detailed procedure of cell maintenance and differentiation is described in detail elsewhere^11^. Briefly, ESCs were plated and passed on feeder cells (mouse embryonic fibroblasts) for 5 days in DMEM-Knockout medium (Gibco, 10829-018) supplemented with 15% fetal bovine serum (Gibco, #10270-42Q4972K), 1% L-Glutamine (Gibco, 25-030-024), 1% Penicillin-Streptomycin (Gibco, 15140-122), 1% non-essential aminoacids (Gibco, 11140-035), 0.24% of 50mM β-mercaptoethanol (Gibco, 31350-010) and 0.0024% of 1mg/ml LIF (EMBL-Heidelberg). After this, cells were passaged on gelatin for two days to dilute feeders out. One gelatin passage was done in DMEM-ES, and the other in IMDM-ES made with IMDM medium (Lonza, BE12-726F).

After the gelatin passages, we plated the cells in a medium for embryoid bodies (EBs) at 0.3×10^6^ cells/10cm petri dish (Sterilin). The EB medium was made of IMDM (Lonza, BE12-726F), 15% fetal bovine serum (Gibco, #10270-42G9552K), 1% Penicillin-Streptomycin (Gibco, 15140-122), 1% L-Glutamine, 0.6% holotransferrin (Roche, 10652202001), 0.0039% MTG (Sigma, M6145) and 50μg/ml ascorbic acid (Sigma, A4544). After 3-3.25 days of EB culture, Flk1^+^ cells were sorted from EBs using rat anti-mouse Flk1-APC antibody (eBioscience, #17-5821-81) with magnetic anti-APC MACS beads (Miltenyi Biotec, #130-090-855). Following the sort, the purified Flk1^+^ cells (> 95% purity) were frozen and stored in liquid nitrogen for further experiments.

### Hemangioblast cultures and experiments

Flk1^+^ cells were plated on gelatin-coated 6-well plates at 1×10^6^ cells/plate in a blast-mix medium containing IMDM (Lonza BE12-726F), 10% fetal bovine EB serum (Gibco, #10270-42G9552K), 0.6% holotransferrin (Roche, 10652202001), 0.0039% MTG (Sigma, M6145), 25μg/ml ascorbic acid (Sigma, A4544), 15% D4T supernatant (EMBL Rome), 0.05% of 10μg/ml VEGF (Peprotech, 100-20) and 0.1% of 10μg/ml IL6 (Peprotech, 216-16). At 24 hours after plating, cells were supplemented either with nothing, 50μM deferoxamine (DFO, Sigma, D9533), 200μM ferric ammonium citrate (Sigma, F5879), either alone or in combination with DFO 50μM. The stock solutions of DFO and ferric ammonium citrate were prepared in sterile distilled water (Gibco) and were added to the cells at a volume, which would not exceed 2% of the total well volume. In some cases, we used a well supplemented with 1-2% water as a control, which was similar to a well with no addition at all (Shvartsman and Lancrin unpublished observations). The time course of hematopoiesis in culture was followed by IncuCyte (Essen Bioscience) time-lapse microscopy from the moment of treatment until 24 hours later. After 24 hours of treatment (unless indicated otherwise), cells were harvested with TrypLE, collected into IMDM with 10% FBS and stained for flow cytometry with one of the following combinations of rat anti-mouse antibodies from eBioscience: CD144-efluor660 (# 50-1441-82) and CD41-PE (# 12-0411-82); CD117-APC (# 17-1171-81) and CD41-PE; any of the combinations above plus CD71-biotin (BDPharmingen, #557416) and streptavidin-AlexaFluor488 (Invitrogen, S11223). The antibody dilutions were 1:200 for CD144-efluor660, CD117-APC and CD71-biotin, 1:400 for CD41-PE. Streptavidin-AF488 was diluted 1:500. Dead cells were excluded by 7-AminoactinomycinD (7AAD, Sigma #A9400) 1:100 staining. Fluorescence was measured with the aid of FACSCanto flow cytometer (BD).

### Cell proliferation assay

Cell proliferation was measured with the Click-It Plus EdU Alexa Fluor 488 flow cytometry assay kit from Invitrogen (C10633), according to manufacturer’s instructions. Briefly, our blast cultures received 10μM EdU for 1 hour before cell harvesting and staining with above-mentioned antibodies. Stained cells were fixed and permeabilized and the ClickIt reaction was performed by manufacturer’s instructions. Fluorescence was measured on FACS-Canto (BD).

### Apoptosis assay

Apoptosis was measured by AnnexinV-FITC apoptosis detection kit (eBioscience #88 8005 72) according to manufacturer’s instructions. Briefly, cells were harvested, stained with the anti-mouse cKit-APC and CD41-PE antibody mix, washed, incubated with Annexin-V-AF488 in AnnexinV-binding buffer, washed and put on ice. 7-AminoactinomycinD was used as a nuclear stain to label dead cells^22^. Fluorescence was measured on FACS-Canto (BD).

### Hematopoietic Colony Forming Unit (CFU) assay

Hemangioblast cultures were grown as described above, but in 20-cm gelatinized dishes (Corning #10314601) instead of 6-well plates. 24 hours after plating, cells were treated or not with DFO 50 μM for 24 hours, then harvested and stained with a mix of anti-mouse CD144-efluor660, CD117-APC and CD41-PE antibodies as described above. Hematopoietic progenitors were gated as VE-Cad^-^ CD41^+^ and sorted into separate sterile falcons according to their Kit/CD41 fluorescence profile (Kit^+^ CD41^+^ or Kit^-^ CD41^+^) with the FACSAria cell sorter. Dead cells were excluded by 7-AAD staining. The cells were plated into CFU mix with 55% methylcellulose, as previously described^3^. Colony number was counted 1 week later. The experiment was independently repeated 5 times.

### Single-cell quantitative RT-PCR

Hemangioblast cultures were grown as above and treated with either nothing or DFO for 24 hours. Cells were harvested and stained with the same combination of anti-mouse antibodies used for colony assays. Single cells from each cell type (Kit^-^ CD41^-^, Kit^+^ CD41^-^, Kit^+^ CD41^+^ and Kit^-^ CD41^+^) were sorted with the FACSAria fluorescence cell sorter (BD Biosciences) into 96-well plates filled with 2 x lysis buffer from the CellsDirect One Step qRT-PCR Kit (Invitrogen, # 11753100) and snap-frozen on dry ice. RT/Specific target amplification reaction (RT-STA) was performed according to manufacturer’s instructions. After RT-STA, individual single-cell qPCR was run on the Fluidigm Biomark HD system. Analysis was performed using R as previously described^11^.

### Labile iron measurements

The method for labile iron measurements with calcein green was adapted from Shvartsman et al^20^. Briefly, cells were stained with calcein-green AM 0.03 μM for 10min at 37 °C, washed, stained with antibodies as described above and sampled by flow cytometry in 10min intervals before (baseline) and after addition of a cell permeable chelator (L1 50 μM). The relative labile iron content is measured as the normalized delta fluorescence (ΔF) between baseline and chelator addition^20^.

### Data analysis and statistics

Flow cytometry raw data were analyzed by exclusion of debris and doublets, exclusion of dead 7AAD+ cells, and gating on different types of live cells according to their VE-Cad/CD41 or Kit/CD41 fluorescence. Single-stained controls were used for compensation. Flow cytometry data were analyzed using the FlowJo 10.2 software (TreeStar Inc.).

Data from flow cytometry experiments were plotted either as frequency (% of cells of a given cell type from total number of cells), or as absolute cell number (the frequency x total cell number (x10^4^)/well calculated by hemocytometer/100). Data were calculated as mean +/- standard deviation or standard error. Each experiment was independently performed from n=3 to n=15. Statistical analysis was either one-way ANOVA followed by Tukey-Kramer multiple comparisons test to compare multiple experimental groups, or paired two-tailed t-test to compare only control versus DFO. Graphs and statistical analysis were done by JMP Pro 12.1.0.

## Acknowledgements

M.S. was supported by the EMBL Interdisciplinary Postdocs (EIPOD) Initiative. The authors thank Kerstin Ganter (EMBL Rome) for excellent technical support, Morgan Oatley (EMBL Rome) for single cell qPCR assistance, Cora Chadick (EMBL Rome) for flow cytometry assistance, Andreas Buness (EMBL Rome), Nicolas Descostes (EMBL Rome) and Bernd Klaus (EMBL Heidelberg) for statistical advice, and Bianka Baying (EMBL Genomics Core Facility) for most valuable support of the sc-q-RT-PCR Fluidigm platform.

## Contributions

M.S. and C.L. conceived the project, M.S. performed experiments, analyzed the data and wrote the manuscript, S.B. performed experiments and analyzed data, C.L. analyzed the data and wrote the manuscript.

## References

1. Medvinsky A, Rybtsov S, Taoudi S. Embryonic origin of the adult hematopoietic system: advances and questions. Development 2011; 138:1017-1031.

2. Al-Drees MA, Yeo JH, Boumelhem BB, Antas VI, Brigden KWL, Colonne CK, Fraser ST. Making blood: The haematopoietic niche throughout ontogeny. Stem Cells International 2015, 2015:571893.

3. Lancrin C, Sroczynska P, Stephenson C, Allen T, Kouskoff V, Lacaud G. The haemangioblast generates haematopoietic cells through a haemogenic endothelium stage. Nature 2009; 457(7231): 892-895.

4. Eilken HM, Nishikawa S, Schroeder T. Continuous single-cell imaging of blood generation from haemogenic endothelium. Nature. 2009; 457(7231): 896-900.

5. Boisset JC, van Cappellen W, Andrieu-Soler C, Galjart N, Dzierzak E, Robin C. In vivo imaging of haematopoietic cells emerging from the mouse aortic endothelium. Nature 2010; 464(7285): 116-120.

6. Batsivari A, Rybtsov S, Souilhol C, Binagui-Casas A, Hills D, Zhao S, Travers P, Medvinsky A. Understanding Hematopoietic Stem Cell Development through Functional Correlation of Their Proliferative Status with the Intra-aortic Cluster Architecture. Stem Cell Reports. 2017;8(6):1549-1562.

7. Boisset JC, Clapes T, Anna Klaus A, Papazian, N, Onderwater J, Mommaas-Kienhuis M, Cupedo T, Robin C Progressive maturation toward hematopoietic stem cells in the mouse embryo aorta. Blood 2015; 125(3): 465–469.

8. McGrath KE, Frame JM, Fegan KH, Bowen JR, Conway SJ, Catherman SC, Kingsley PD, Koniski AD, Palis J. Distinct sources of hematopoietic progenitors emerge before HSCs and provide functional blood cells in the mammalian embryo. Cell Rep. 2015;11(12):1892-1904.

9. Kang HJ, Mesquitta WT, Jung HS, Moskvin OV, Thomson JA, Slukvin II. GATA2 is dispensable for specification of hemogenic endothelium but promotes endothelial-to-hematopoietic transition. Stem Cell Reports. 2018; 11(1): 197–211.

10. Vargel Ö, Zhang Y, Kosim K, Ganter K, Foehr S, Mardenborough Y, Shvartsman M, Enright AJ, Krijgsveld J, Lancrin C. Activation of the TGFβ pathway impairs endothelial to haematopoietic transition. Scientific Reports 2016; 6:21518.

11. Bergiers I, Andrews T, Vargel Bölükbaşı Ö, Buness A, anosz E, Lopez-Anguita N, Ganter K, Kosim K, Celen C, Itır Perçin G, Collier P, Baying B, Benes V, Hemberg M, Lancrin C. Single-cell transcriptomics reveals a new dynamical function of transcription factors during embryonic hematopoiesis. Elife 2018; 7 pii: e29312.

12. Pereira CF, Chang B, Gomes A, Bernitz J, Papatsenko D, Niu X, Swiers G, Azzoni E, de Bruijn MF, Schaniel C, Lemischka IR, Moore KA. Hematopoietic reprogramming in vitro informs in vivo identification of hemogenic precursors to definitive hematopoietic stem cells. Developmental Cell 2016; 36(5): 525-539.

13. Muckenthaler MU, Rivella S, Hentze MW, Galy B. A red carpet for iron metabolism. Cell 2017;168(3): 344-361.

14. Levy JE, Jin O, Fujiwara Y, Kuo F, Andrews NC. Transferrin receptor is necessary for development of erythrocytes and the nervous system. Nature Genetics 1999; 21(4):396-9.

15. Gunshin H, Fujiwara Y, Custodio AO, DiRenzo C, Robine S, Andrews NC. Slc11a2 is required for intestinal iron absorption and erythropoiesis but dispensable in placenta and liver. J Clin Invest. 2005 May;115(5):1258-66.

16. Mihaila C, Schramm J, Strathmann FG, Lee DL, Gelein RM, Luebke AE, Mayer-Proschel M. Identifying a window of vulnerability during fetal development in a maternal iron restriction model. PLoS ONE 2011; 6(3): e17483.

17. Haider BA, Olofin I, Wang M, Spiegelman D, Ezzati M, Fawzi WW. Anaemia, prenatal iron use, and risk of adverse pregnancy outcomes: systematic review and meta-analysis. British Medical Journal 2013;346:f3443

18. Macedo MF, De Sousa M, Ned RM, Mascarenhas C, Andrews NC, Correia-Neves M. Transferrin is required for early T-cell differentiation. Immunology 2004; 112: 543-549.

19. Ned RM, Swat W and Andrews NC. Transferrin receptor 1 is differentially required in lymphocyte development. Blood 2003; 102(10): 3711-3718.

20. Shvartsman M, Fibach E, Cabantchik ZI. Transferrin-iron routing to the cytosol and mitochondria as studied by live and real-time fluorescence. Biochemical Journal 2010; 429: 185-193.

21. Iacovino M, Bosnakovski D, Fey H, Rux D, Bajwa G, Mahen E, Mitanoska A, Xu Z, Kyba M. Inducible cassette exchange: a rapid and efficient system enabling conditional gene expression in embryonic stem and primary cells. Stem Cells. 2011; 29(10):1580-8.

22. Vermes I, Haanen C, Steffens-Nakken H, Reutelingsperger C. A novel assay for apoptosis. Flow cytometric detection of phosphatidylserine expression on early apoptotic cells using fluorescein labeled Annexin V. Journal of Immunological Methods 1995; 184: 39-51.

23. Chen Z, Zhang D, Yue F, Zheng M, Kovacevic Z, Richardson DR. The iron chelators Dp44mT and DFO inhibit TGF-β-induced Epithelial-Mesenchymal Transition via Up-Regulation of N-Myc Downstream-regulated Gene 1 (NDRG1). J Biol Chem. 2012;287(21):17016-17028.

24. Pearson S, Cuvertino S, Fleury M, Lacaud G, Kouskoff V. In vivo repopulating activity emerges at the onset of hematopoietic specification during embryonic stem cell differentiation. Stem Cell Reports 2015; 4(3): 431–444.

25. Cornock R, Gambling L, Langley-Evans SC, McArdle HJ, McMullen S. The effect of feeding a low iron diet prior to and during gestation on fetal and maternal iron homeostasis in two strains of rat. Reprod Biol Endocrinol. 2013;11:32.

